# Success of *Escherichia coli* O25b:H4 ST131 clade C associated with a decrease in virulence

**DOI:** 10.1101/786350

**Authors:** Marion Duprilot, Alexandra Baron, François Blanquart, Sara Dion, Philippe Lettéron, Saskia-Camille Flament-Simon, Olivier Clermont, Erick Denamur, Marie-Hélène Nicolas-Chanoine

## Abstract

*Escherichia coli* of sequence type (ST) 131 resistant to fluoroquinolones and producer of CTX-M-15 is globally one of the major extraintestinal pathogenic *E. coli* (ExPEC). ST131 phylogenesis showed that multidrug-resistant ST131 strains belong to a clade called C, descending from an ancestral clade called B, comprising mostly antibiotic-susceptible strains. Antibiotic resistance could appear as one of the keys of the clade C global success. We hypothesized that other features of ST131 clade C could contribute to this success since other major global ExPEC clones (ST73, ST95) are mostly antibiotic-susceptible. To test this hypothesis, we measured the growth abilities, early biofilm formation and virulence-factor content of a collection of clade B and clade C strains. Moreover, using competition assays, we measured the capacity of selected representative strains of clades B and C to colonize the mouse intestine and urinary tract, and to kill mice in a septicemia model. Clade B and C strains had similar growth ability. However, clade B strains were more frequently early biofilm producers, expressed mostly faster their type 1 fimbriae and displayed more virulence factor-encoding genes than clade C strains. Clade B outcompeted clade C in the gut and/or urinary tract colonization models and in the septicemia model. These results strongly suggest that clade C strain evolution includes a loss of virulence, *i.e*. a process that could enhance micro-organism persistence in a given host and thus optimize transmission. This process, associated with acquired antibiotic-resistance, could ensure clade C strain survival in environments under antibiotic pressure.

**Importance:** Extraintestinal pathogen *Escherichia coli* (ExPEC) are virulent but mostly antibiotic-susceptible. One worrying exception is ST131, a major multidrug resistant ExPEC clone that has spread worldwide since the 2000s. To contain the emergence of this threatening clone, we need to understand what factors favored its emergence and dissemination. Here, we investigated whether multidrug-resistant ST131 had advantageous phenotypic properties beyond multidrug resistance. To this end, we competed the emergent multidrug-resistant ST131 with its antibiotic-susceptible ancestor in different conditions: biofilm production, *in vivo* colonization and virulence experiments. In all *in vivo* competitions, we found that multidrug-resistant ST131 was losing to its ancestor, suggesting a lesser virulence of multidrug-resistant ST131. It was previously described that losing virulence can increase micro-organism persistence in some populations and subsequently its level of transmissibility. Thus, a decreased level of virulence, associated with multidrug resistance, could explain the global success of ST131.

## Introduction

In the last two decades, *Escherichia coli* O25b:H4 of sequence type (ST) 131 has emerged worldwide among human extraintestinal pathogenic *E. coli* (ExPEC) (1, 2). This multidrug resistant clone has become a major public health issue. The evolutionary history of this successful ST has been reconstructed in details. Within O25b:H4 ST131, two clades called B and C were distinguished on the basis of their *fimH* gene allele, *H*22 and *H*30, respectively (3, 4). Through a time-calibrated phylogeny, Ben Zakour *et al.* showed that clade B is the ancestor of clade C and that sequential mutational events have shaped the evolution of ST131 from the 1950s (5). This evolutionary scenario shows the diversification of clade B into different subclades (from B1 to B5, then the intermediate B0) and notably clade C. Clade C also diversified into different subclades characterized by the acquisition of the *fimH*30 allele around 1980 (subclade C0), followed by the acquisition of the *gyrA*-1AB and *parC*-1aAB alleles encoding fluoroquinolone resistance that occurred around 1987 (subclades C1 and C2). All subclade C2 strains and some of subclade C1 forming C1-M27 cluster (6) are additionally resistant to extended spectrum cephalosporins (ESC) due to the production of extended spectrum β-lactamases (ESBL) of CTX-M type, CTX-M-15 and CTX-M-27, respectively. Epidemiologic studies mostly determined ST131 prevalence among *E. coli* isolates resistant to fluoroquinolones and/or producers of ESBL (7). Studies designed to assess the relative frequencies of clade B and clade C ST131 strains are rare (8–10) and those to assess the relative frequencies of B and C subclades strains do not exist. Recently, Kallonen *et al.* depicted 14.4% of ST131 clades among bacteremia *E. coli* isolates systematically collected between 2001 and 2012 in England (9). Using the English ST131 isolates’ whole genome sequences, we were able to assess the relative frequencies of subclades B and C in the present study.

The evolutionary success of ST131 is still largely unexplained. Resistance to antibiotics, notably to fluoroquinolones and ESC may explain the success of clade C strains, as these resistances were shown not to impact the clade C growth fitness (11, 12). However, clones susceptible to antibiotics, such as ST73 and ST95, are as successful as ST131 clade C among bacteremia, suggesting that factors other than antibiotic resistance can participate in the success of a clone (9), The aim of our study was thus to investigate the potential role of factors other than antibiotic resistance, notably growth rate, biofilm formation, colonization ability and virulence, in the success of ST131 clade C strains. To this purpose, we first established a collection of 39 *E. coli* O25b:H4 ST131 comprising representatives of the ST131 clade and subclade genetic diversity and tested them in a variety of *in vitro* phenotypic tests. We then selected three strains for further *in vitro* and *in vivo* competition assays, including various mouse models.

## Materials and methods

Technical details for each section are available in the supplemental material Text S1.

### Bacterial strains

The 39 studied O25b:H4 ST131 *E. coli* strains comprised 18 *fimH*22 and 21 *fimH*30 strains obtained between 1993 and 2012 from different geographic origins and sources (Table S1). Nalidixic acid and ciprofloxacin resistance had been determined by the agar diffusion method and interpreted following the 2015 EUCAST recommendations (www.eucast.org) and ESBL production by the double disk synergy test (13). CFT073 and *E. coli* K-12 MG1655 strains were used as positive and negative controls, respectively, in the septicemia mouse model.

### Genome sequencing and analysis

Whole genome sequencing (WGS) of our 39 strains was performed (Tables S2). All genomes were analyzed for plasmid content and typing [Plasmid Finder, with identity >95%, and pMLST (14)]. Then, Abricate (15) was used to detect genes encoding antibiotic resistance with Resfinder (16) and virulence factors with a custom virulence database composed of VirulenceFinder (17), the virulence factor database (VFDB) (18), and classical genes characterizing ExPEC. Virotypes were determined as previously described (19). All contigs were submitted to the MicroScope Platform (20) for further gene investigation such as *gyrA/parC* alleles and genes involved in the biofilm formation (21). When necessary, presence or absence of some genes was controlled by PCR.

We investigated how the different subclades B and C of ST131 were represented in our collection and how they evolved in frequency over time in a larger collection from Kallonen *et al.* (9). To that end, we complemented our 39 genomes with 218+21 genomes from two published studies [218 from Kallonen *et al.* (9), 21 from Ben Zakour *et al.* (5)]. The phylogenetic tree including all the strains was constructed from non-recombinant single nucleotide polymorphisms (SNPs) of core genomes using maximum likelihood. To infer the linear trend of Kallonen *et al.*’s isolates over 2001-2012, we fitted a logistic model to the frequency of each subclade as a function of time (in years).

### Gene deletion and gene complementation

Replacement by a kanamycin resistance cassette was used to inactivate the chromosomal *fimB* gene of S250 and CES131C strains and the plasmidic *aadA2* gene of CES131C strain following a strategy adapted from Datsenko and Wanner (22). Primers and plasmids used are listed in Tables S3 and S4. Complementation of Δ*fimB* mutants were performed by cloning the promoter and encoding regions of the parental *fimB* genes into pSC-A-amp/kan (Table S4) by using the StrataClone PCR Cloning Kit (Agilent Technologies, Massy, France). The recombinant plasmid was then electroporated into competent strains S250Δ*fimB*:: or CES131CΔ*fimB*:: and transformants were selected on lysogeny broth (LB) (Invitrogen, Carlsbad, California, USA) agar plates containing 100 mg/L of kanamycin. The empty plasmid pSC-A-amp/kan-Control_Insert was used as negative control. All mutants were confirmed by PCR and sequencing.

### Determination of early biofilm formation

The primary step (5 h incubation) of biofilm formation was measured using BioFilm Ring Test^®^ (BioFilm Control, Saint-Beauzire, France) according to the manufacturer’s recommendations and as previously described (23). The biofilm formation index (BFI), with values ranging from 20 (absence of biofilm formation) to 0 (high biofilm formation), is inversely proportional to the biofilm formation ability. A BFI value of 10 was chosen as the biofilm production cut-off (biofilm formation: BFI≤10, no biofilm formation: BFI>10). The experiments were repeated at least three times for each strain. BHI broth without any strains was used as negative control, and strains S250 and 39, previously described with this method as strong and never biofilm producers, respectively, as control strains (24).

### Yeast cell agglutination assay

Expression of type 1 fimbriae was assessed by using the yeast cell (*Saccharomyces cerevisiae*) agglutination assay as previously described (25), and after adaptations to highlight the early expression of type 1 fimbriae, by using 10 µL of a pellet obtained after centrifugation (3000 g, 10 min) of 3 mL of LB broth culture after 2 and 5 h-incubations for the agglutination assay. Based on the biofilm formation and the sequence of the *fim* operon, strains S250 and H1447 were used as positive and negative controls, respectively.

### Congo red assays

Curli and cellulose production was assessed by using stationary-phase cultures spotted onto LB low salt (Invitrogen) plates supplemented with 40 µg/mL Congo red (Sigma Aldrich, St-Quentin Fallavier, France) at 30°C during 48 h, as previously described (26). The production of curli fimbriae and cellulose resulted in the red, dry and rough (rdar) colony morphotype.

### Colicin and/or phage production

Colicin and/or phage production was detected by plaque lysis assays, with *E. coli* K-12 used as the sensitive strain as previously described (27). The assay was considered as positive when tested strains were surrounded by a halo, corresponding to the growth inhibition of *E. coli* K-12 strain.

### Fitness assays

#### Individual fitness assays

Fitness assay was performed as previously described in LB broth (28) by using an automatic spectrophotometer (Tecan Infinite F200 Pro) that measures the OD_600_ in each well every 5 min over a period of 24 h. The experiment was repeated three times. Growth curves were then analyzed and maximum growth rates was calculated and expressed in h^-1^.

#### Competition experiments

Selected strains of our collection and constructed mutants were submitted in pairs to competition in different conditions.

##### *In vitro* planktonic competitions

To determine the relative fitness of strains in planktonic conditions, they were grown in couple in LB broth during 100 generations, *i.e.* during ten days, as previously described (29). Competitive indexes (CI) were obtained using the following formula: logCFU[(isolate1/isolate2)T_x_/(isolate1/isolate2)T_0_], isolate1 being the first strain cited and isolate2 the second one in figure legends.

##### *In vitro* competitions in biofilm conditions

To determine the relative fitness of strains in biofilm conditions, they were grown at 1:1 ratio in mixed colonies during seven days in aerobic and anaerobic conditions, with adaptations to what’s previously described (30). For each competition, the experience was repeated three times. CIs were obtained as described above.

##### Intestinal colonization mouse model

Six-week-old female mice CD-1 from Charles River® (L’Arbresle, France) pre-treated with streptomycin before inoculation of challenging strains were used to assess strains’ relative ability to colonize the mouse intestine, as previously described (28). At least five mice were used for each competition. At days 1, 4 and 7, the intestinal population of *E. coli* was estimated by plating dilutions of weighed fresh feces on LB agar and LB agar with appropriate antibiotics (kanamycin 50 mg/L or ciprofloxacin 1 mg/L). CIs were obtained as described above.

##### Septicemia mouse model

Female mice OF1 of 14-16 g (4-week-old) from Charles River^®^ were used to assess the strains’ relative ability to cause sepsis, as previously described (31). After subcutaneous inoculation of individual strain or mixed strains at a 1:1 ratio, time to death was monitored during seven days. Five to ten mice were used for each assay. In all experiments, CFT073 strain and *E. coli* K-12 MG1655 strain were used as positive and negative control, respectively (32). Kaplan-Meier estimates of mouse survival were performed for individual strain inoculations. In competitions assays, the spleen of all spontaneously dead mice was collected, weighed and pounded in physiological water and infecting population of *E. coli* was estimated by plating dilutions of spleen suspensions on LB agar and LB agar with appropriate antibiotics (kanamycin 50 mg/L, ciprofloxacin 1 mg/L or ampicillin 100 mg/L). CIs were obtained as described above.

##### Urinary tract infection mouse model

CBA female mice of 8-22g (8-week-old) from Janvier^®^ (Le Genest-Saint-Isle, France) were used to assess strains’ relative ability to cause an ascending unobstructed urinary tract infection, as previously developed in our lab (33). Ten mice were used for each competition assay, and the experiments were repeated twice. CIs were obtained as described above.

### Statistical analysis

Wilcoxon rank sum test was used to compare the average number of virulence factor (VF) per clade and Fisher’s exact test was used to test the distribution of each VF-encoding gene between clades. Individual fitness measurement was estimated by a mixed model with random effect on the strain to take into account the triplicate determination. For biofilm data, two-way repeated measures ANOVA was performed, followed by Tukey’s range test when necessary (if there were three groups). The association between groups (virotypes, biofilm phenotypes, yeast agglutination test and Congo red assay) was assessed by Fisher’s exact test. For the *in vitro* and *in vivo* competitions, a non-parametric Wilcoxon test on paired data was conducted on CI values, and P values were corrected for multiple comparisons by the Benjamini-Hochberg procedure (34) when necessary. Mouse survival differences were determined by Log-Rank test. A significance level of 0.05 was used for all tests. All statistical analyses were carried out with R software (35).

To visualize all data of our strain collection in one graph, a multiple correspondence analysis (MCA) was realized with R software (35). The variables included in the analysis were strain source, *fimH* allele, fluoroquinolone susceptibility, ESBL production, and biofilm formation phenotype. The distance between the variables and the origin of the graph measures the quality of their representation and is estimated by the square cosine (cos2): variables far from the origin of the graph are discriminant and got a high cos2 and variables close to the origin are not perfectly depicted and got a low cos2 (Figure S1A).

### Ethics statement

Colonization, septicemia and pyelonephritis murine protocols (n°APAFIS#4948-2016021216251548 v4, n°APAFIS#4951-2016020515004032 v2 and n°APAFIS#4950-2016021211417682 v4, respectively) were approved by the French Ministry of Research and by the ethical committee for animal experiments.

### Data availability

Raw sequences of the 39 isolates were deposited in GenBank under BioProject accession number <u>PRJNA320043</u> and <u>PRJNA566165</u>.

## Results

### Evolutionary success of the C1 and C2 subclade strains in extraintestinal infections

Using both the genome sequences of the ST131 strains published by Kallonen *et al.* (9) and the genome sequences of strains belonging to the different B and C subclades (5), we characterized the B and C subclades of Kallonen *et al.*’s O25b:H4 ST131 strains (Figure S2). By investigating trends in O25b:H4 ST131 subclade frequency over the period 2001-2012 (Figures 1A, 1B, 1C) in extraintestinal infections, we observed that (i) B4 and B5 subclade strains increased more than the other B subclade strains, (ii) clade C strains increased more than clade B strains and (iii) C2 subclade strains displayed the most important increase. Thus, although both clades B and C were maintained over this time period, clade C appears more evolutionarily successful than clade B.

**Figure 1.**
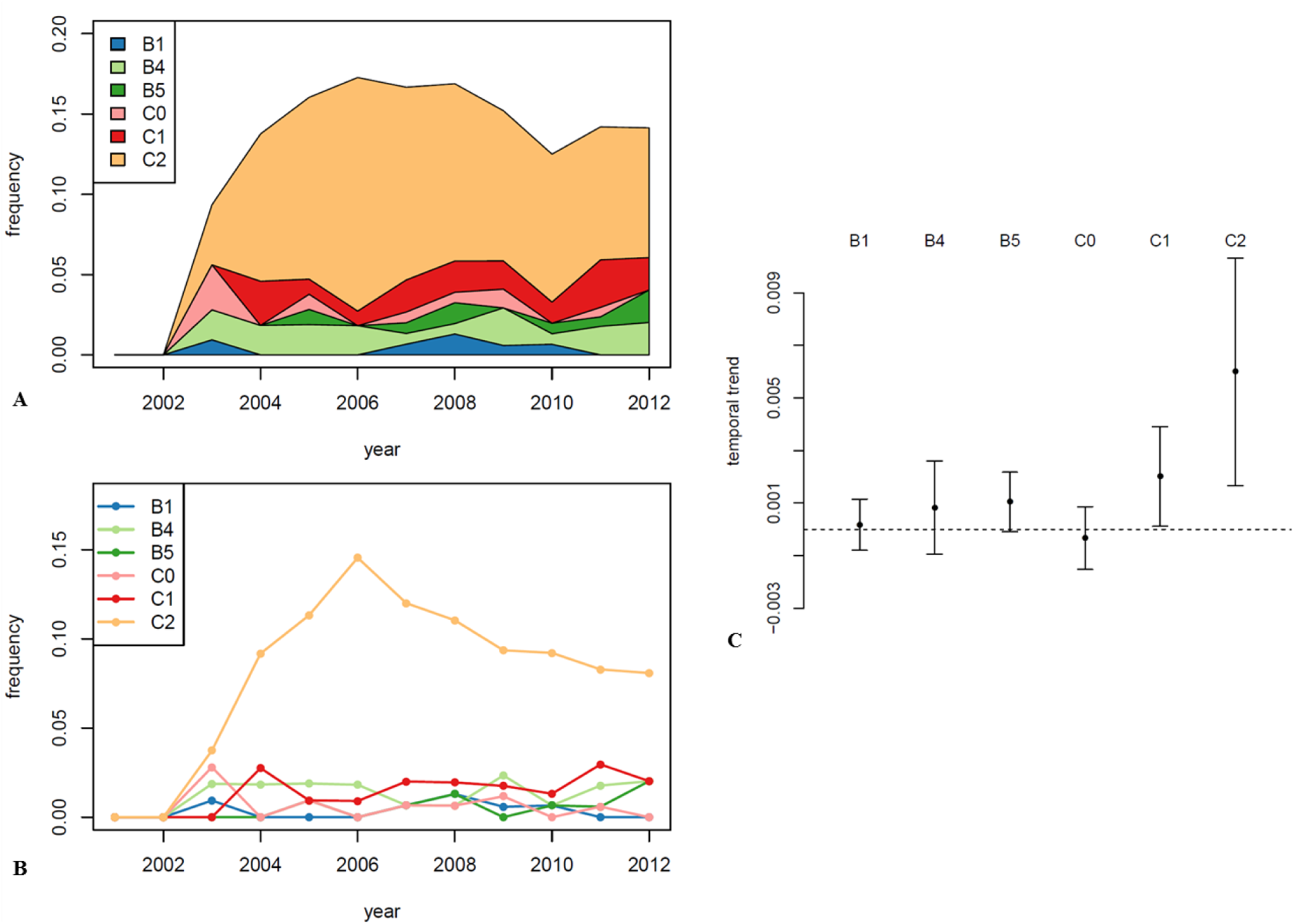
Dynamics of subclades of O25b:H4 ST131 *E. coli* isolated from bacteremia over a 11-year sampling period [data from Kallonen *et al* (9)]. A. Cumulated frequencies of ST131 subclades as a function of time, from the most ancestral subclade B1 at the bottom to the derived subclade C2 on top. B. Frequency of ST131 subclades as a function of time. Subclades are represented by colors. C. Inferred linear trends of ST131 subclade frequencies (among all *E. coli* strains) as a function of time over 2001-2012.

### Establishment of a collection of representative strains of the ST131 O25b clades and subclades

#### Clades and subclades

To determine which previously established ST131 clades and subclades our 39 strains belonged to, a phylogenetic tree was inferred using core genomes exempt of recombinant regions (Figure S2). Our collection comprised strains belonging to each subclade B (except for B2 and B0) and each subclade C. Among the 19 clade B strains, 17 displayed the expected *fimH*22, one a fim*H22*-like variant, and one, CES131C strain belonging to subclade B4, the *fimH*30 variant that had been described as specific of clade C strains. For these reasons, CES131C strain was called “Hybrid” in the rest of the study.

#### Plasmids

All strains but one (strain 208) harbored an IncF plasmid with a FII, FIA and FIB replicon allele composition corresponding to the one previously described for clade B and C1 and C2 subclades strains (12), except that one C1 subclade strain (CES9C) lacked the FIB replicon as described in C2 subclade strains (Table S5).

#### Genes encoding antibiotic resistance

As shown in Table S5, our collection comprised strains that displayed a diverse antibiotic resistance-encoding gene content in both clades B and C, with a gradual accumulation of these genes from clade B to clade C.

#### VF-encoding genes

Among the genes classically sought for in ExPEC (19), those found in our strains are presented according to subclades in Table S6. The average number of VF-encoding genes was higher in clade B strains (17±2.1) than in clade C strains (14.3±1.3) (P<0.0001). The *hlyF, cdtB*, *iroN, kpsMII*, *cvaC, iss* and *ibeA* genes were significantly more frequent among clade B strains than among clade C strains whereas the opposite was found for the *sat* gene. Virotypes (19) were significantly associated with subclades (P<0.00001) and virotypes not yet described were displayed by Hybrid, one C1 subclade and three C2 subclade strains.

### Individual strain phenotypic assays *in vitro*

#### Maximum growth rate (MGR)

We assessed the fitness of our 39 strains by measuring MGR in planktonic conditions (Figure 3). MGR did not differ significantly between clade B and clade C strains (P = 0.08), and between clinical and feces isolates (P=0.07) (Figures 3A and 3B). Within clade B strains, those resistant to nalidixic acid seemed to have a lower MGR than the susceptible ones (Figure 3C), but this difference was not significant (P=0.2). Regarding clade C strains, no significant MGR differences were observed between nalidixic acid/ciprofloxacin-resistant strains and susceptible strains (P=0.9) (Figure 3D).

### Kinetic of early biofilm formation and expression of type 1 and curli fimbriae

Clade B strains were more frequently biofilm producers than clade C strains at 2 and 5 h (P<0.001). We found three significantly different phenotypes of early biofilm production (P<0.0001) within our 39 strains (Figure 2, Table S5): early and persistent producers (BFI≤10 at 2, 3 and 5 h), delayed producers (BFI>10 at 2 and 3 h but ≤10 at 5 h) and never producers (BFI>10 at 2, 3 and 5 h). As BFI values after 2 and 3 h of incubation classified a given strain into the same biofilm production phenotype (Figure 2A), only BFI values obtained after 2 and 5 h of incubation were considered for further analyses. The 20 clade C strains were either delayed producers (n=8) or never producers (n=12). There was no significant association between subclades C and biofilm phenotype. Clade B strains produced biofilm more frequently than clade C strains. Five (71%) of the B5 strains were early producers and two (29%) were delayed producers, whereas all eight B4 strains were delayed or never producers. This difference between B5 and B4 strains was statistically significant (P=0.02).

**Figure 2.**
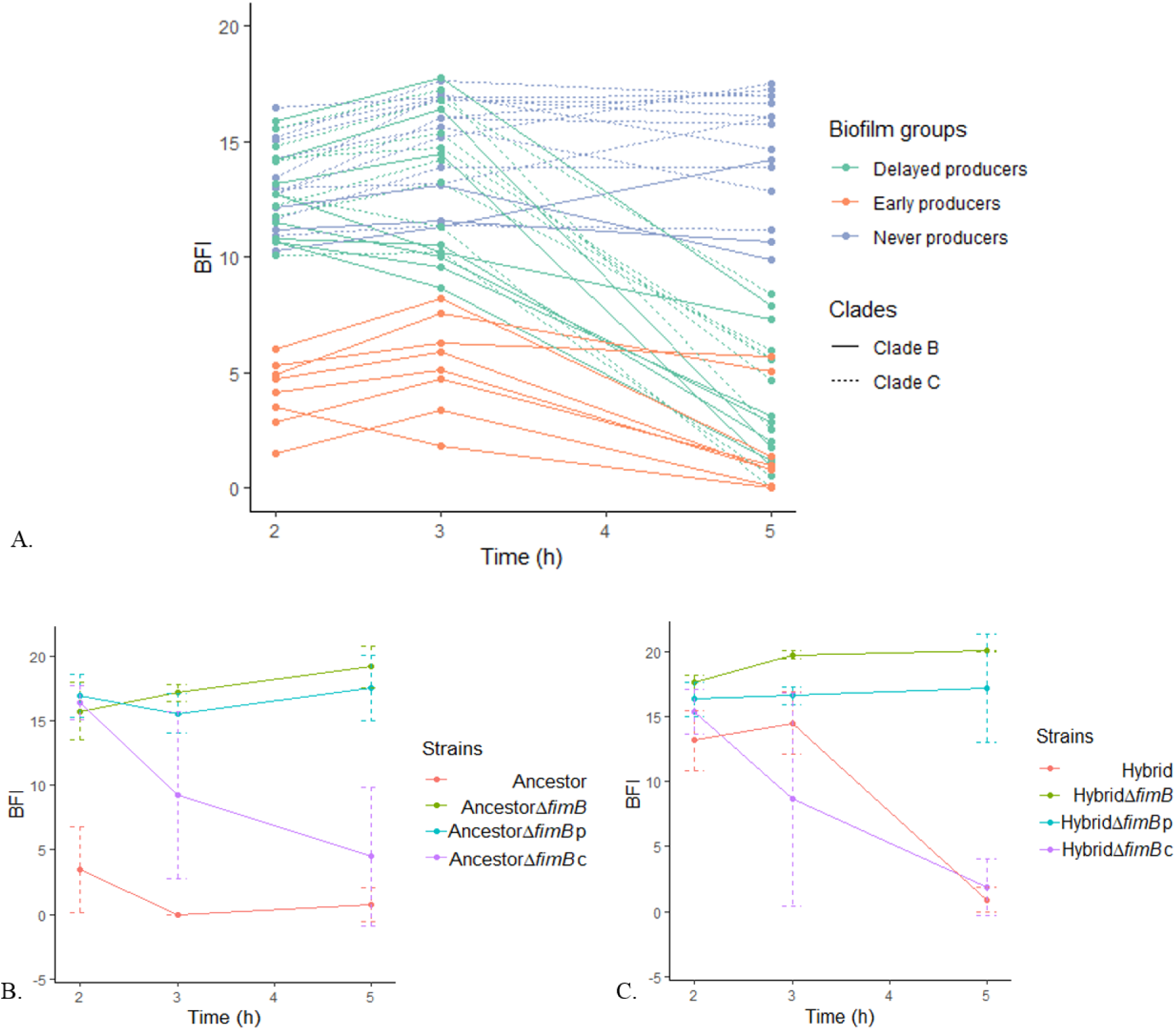
Early biofilm production. Biofilm Ring Test^®^ was used to measure biofilm formation after 2, 3 and 5h of static incubation. BFI values are inversely proportional to the biofilm formation capacity. A. Biofilm formation for the 39 *E. coli* ST131 strains. Clade B: continuous line; clade C: dashed line. Biofilm groups: blue lines: never producers; green lines: delayed producers; orange lines: early producers. B and C. Biofilm measurements for the *fimB* gene knockout experiments on Ancestor and Hybrid strains, respectively. Δ*fimB*: Δ*fimB*::; Δ*fimB*p: Δ*fimB*::pSC-A, strain complemented with an empty plasmid; Δ*fimB*c: Δ*fimB*::pSC-A-*fimB*, strain complemented with its *fimB* gene.

**Figure 3.**
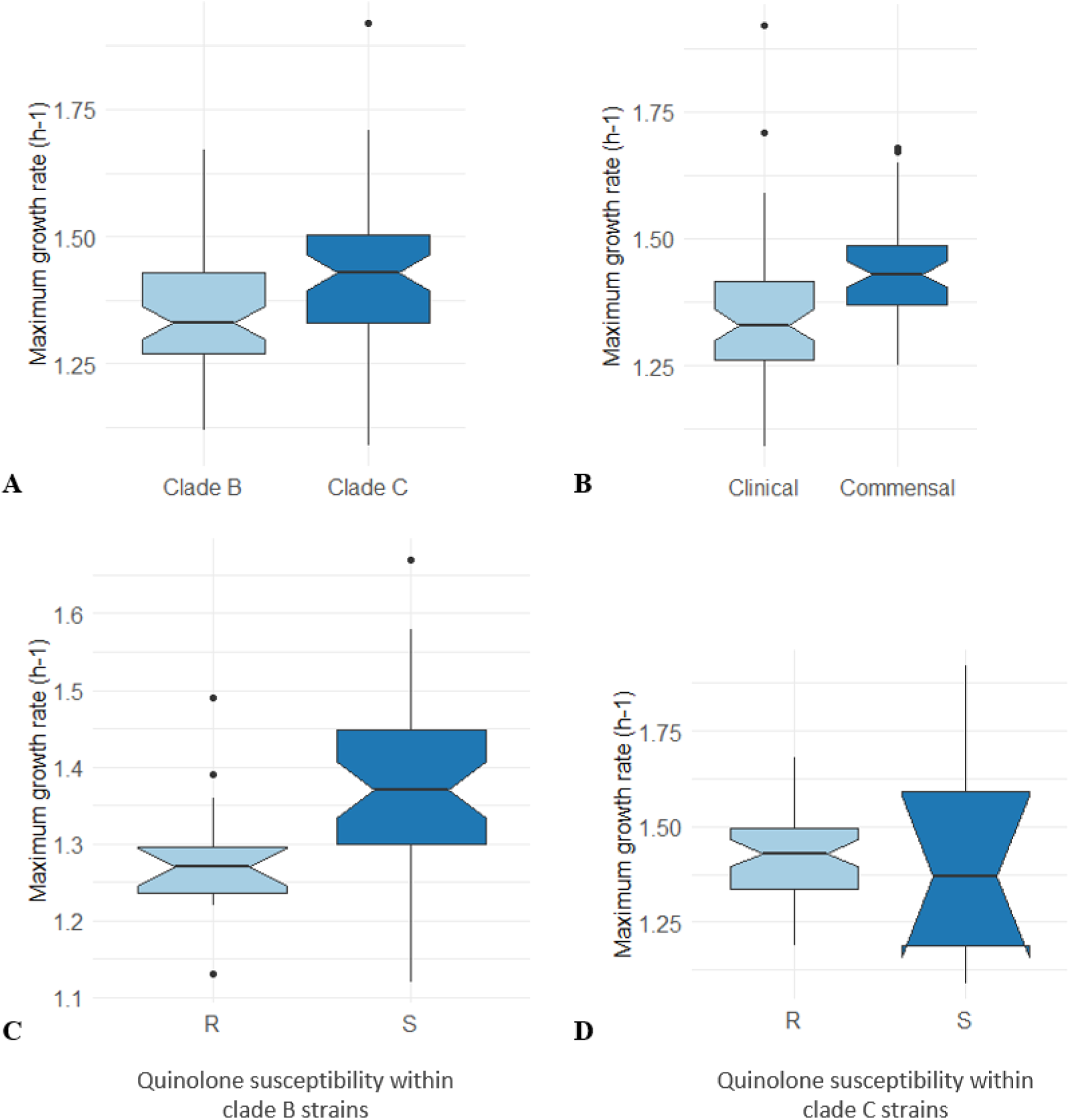
Maximum growth rate of the 39 *E. coli* strains in LB. Maximum growth rate is calculated from three independent culture assays and expressed in h^-1^. Boxplots indicate maximum growth rate distribution across strains, and horizontal black bars indicate median values. The upper and lower ends of the box correspond to the upper and lower quartiles, respectively. Error bars represent the standard error of the mean of three experiments. Outliers are represented by dots. Strains were grouped into two categories: A. Clade B and clade C strains; B. Clinical and commensal (feces) strains; C. Nalidixic acid-resistant (R) and susceptible (S) strains within clade B; D. Nalidixic acid-resistant (R) and susceptible (S) strains within clade C. No significant difference was observed in each comparison (Wilcoxon test)

As the ability to form early biofilm is linked to type 1 fimbriae and curli fimbriae expression (36), we studied these expressions in our strains by using the yeast agglutination test and analyzing colony aspect of bacteria spotted on Congo red agar, respectively. After 2 and 5 h of shaking growth, we observed that the early biofilm producers (n=8) expressed their type 1 fimbriae more quickly than the delayed and never producers (P≤0.0001) (Table 1). Applying the standard yeast agglutination test as Totsika *et al.* (25) in which fimbriae expression is assessed after ≥ 24 h, no significant differences were observed for type 1 fimbriae expression according to biofilm phenotypes or clade membership, in both shaking and static conditions (Table 1). Production of curli fimbriae and cellulose was explored (Table 2). There was a significant association between Congo red morphotype and early biofilm formation capacity (P<0.0001). Significantly more clade B (47%) than clade C strains (15%) were positive for the Congo red assay (P<0.05).

**Table 1.**
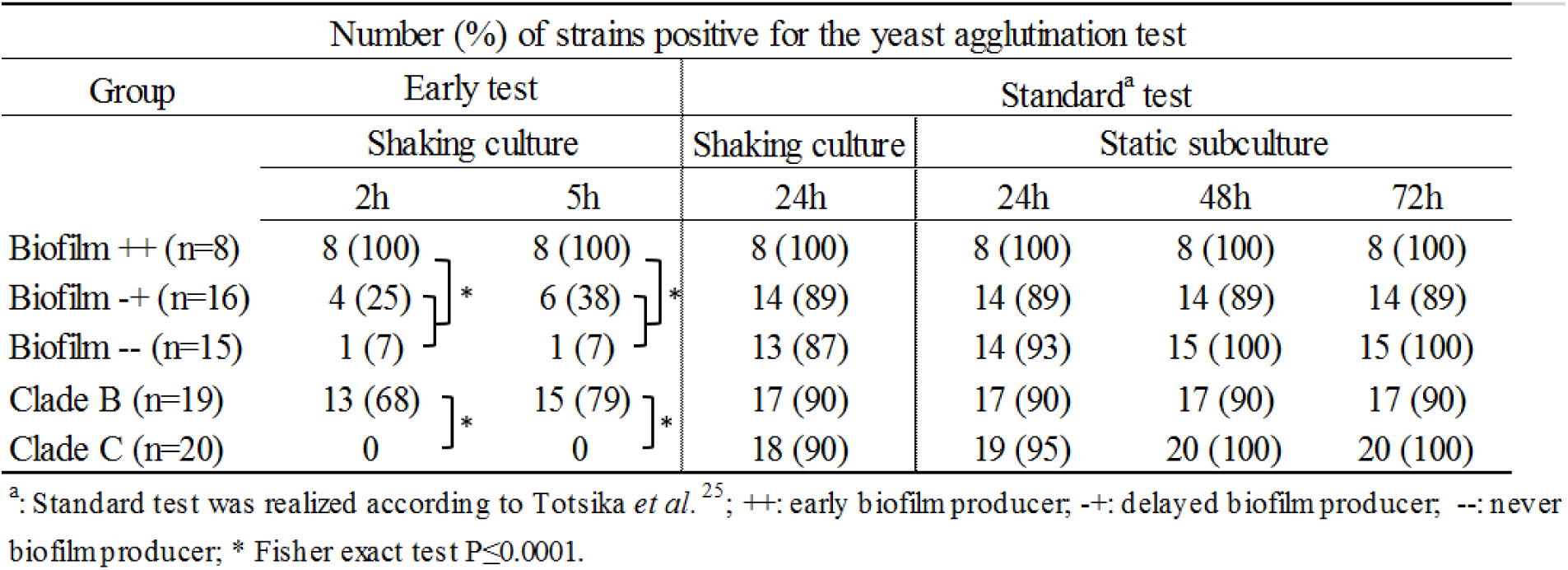
Yeast agglutination test applied to the 39 strains in different growth conditions and at different time points according to biofilm production phenotype and clade type.

**Table 2.** Colony color and morphotype on Congo red agar plates of the 39 strains according to biofilm production phenotype and clade type.

| Number (%) of strains with a positive Congor red test |  |  |
| --- | --- | --- |
| Group | Congo red staining | Rdar morphotype |
| Biofilm ++ (n=8) | 8 (100) | 8 (100) |
| Biofilm -+ (n=16) | 1 (6) | 1 (6) |
| Biofilm -- (n=15) | 4 (27) | 3 (20) |
| Clade B (n=19) | 9 (47) | 9 (47) |
| Clade C (n=20) | 3 (15) | 2 (10) |

++: early biofilm producer; -+: delayed biofilm producer; --: never biofilm producer; Rdar : red, dry and rough colonies; \* Fisher exact test P<0.0001;
\*\* Fisher exact test P<0.001

#### Genes potentially linked to different phenotypes of early biofilm formation

To investigate the genetic factors potentially involved in differences in biofilm formation, we examined the diversity in the *fim* operon across all strains, knowing that Fim proteins participate in biofilm formation (21) and that an insertion sequence ISEc55 has been described in the clade C strain *fimB* gene (25) that encodes a co-factor of the type 1 fimbriae synthesis regulation. We found at least one non-synonymous SNP per gene in the *fim* operon between clade B and clade C strains (Figure S3). The entire *fim* operon was the same in all clade C strains, including ISEc55 within the *fimB* gene. Within clade B, two strains (H1447 and 001-001) found as delayed biofilm producers possessed a partially deleted *fim* operon.

Hybrid, which was also a delayed biofilm producer, displayed a *fim* operon completely different from that of the other clade B strains, showing a perfect match with the *fim* operon of clade C strains, except for two SNPs and the absence of ISEc55 within its *fimB* gene (Figure S3). In order to assess the role of the *fimB* gene inactivation in clade C, we constructed Δ*fimB* variants from an early biofilm producer B1 strain (S250 strain), and a delayed biofilm producer (Hybrid). Both Δ*fimB* variants lost their ability to form biofilm during the 5h-time period (Figures 2B and 2C). Complementation with the wild *fimB* gene restored the biofilm production phenotype displayed by the parental strain of the two variants (Figures 2B and 2C). Δ*fimB* variants were also submitted to yeast agglutination test and Congo red assay. Both Δ*fimB* variants were negative for early yeast agglutination test, unlike parent strains, but were positive for standard test. No changes were observed on Congo red phenotype for both variants (data not shown).

Further genes and/or deduced proteins previously described as involved in *E. coli* biofilm formation (21) were compared between our strains by taking S250 strain as reference (data not shown). We observed identical protein sequence modifications in all B4 strains that were all delayed biofilm producers: substitution K142Q in FlgD that is required for flagellar hook assembly (37), the presence of the *nanC* gene (absent in the other clade B strains) and encoding N-acetylneuraminic acid porin of which regulators also control the *fimB* expression (38), and a 7-bp deletion at the end of the *ydaM* gene encoding a diguanylate cyclase playing a role in the curli biosynthesis by inducing the transcription of the regulator CsgD.

### Strain selection for competition assays

In order to compare more accurately the fitness of clade B and C strains, we turned to competitions assays. To select candidates for competitions among our 39 strains, we used the phylogenetic history (Figure S2), the multiple correspondence analysis (Figure S1B), acquired antibiotic resistance (Table S5) and MGR values. Among clade B strains, we selected strain S250 *i.e*. one of the two most ancestral strains (subclade B1) that we called “Ancestor”.

Among clade C strains, we selected a C1 subclade strain, CES164C, harboring the clade C-characteristic fluoroquinolone resistance and a limited plasmid-mediated resistance (amoxicillin resistance related to TEM enzyme), to avoid the potential burden of additional resistances, such as that related to the production of CTX-M-15 in each C2 subclade strain. Accordingly, we called CES164C strain “Emergent”. We also retained Hybrid (subclade B4) that harbors the first trait having characterized the emergence of clade C strains *i.e.* the *fimH*30 allele.

As shown in Table 3, Ancestor, Emergent and Hybrid displayed a similar average MGR in LB (Wilcoxon Rank Test, P>0.1), were non-colicin/phage producers, but displayed differences in early biofilm formation, type 1 fimbriae expression and Congo red phenotype, as well as in the number and content of genes encoding antibiotic resistance and VFs (Table 3 and Figure 4). However, some VF-encoding genes were present only in Ancestor: the invasin-encoding *ibeA* gene, the salmochelin-encoding *iroBCDEN* cluster, and the *etsC* gene encoding a putative secretion system I, whereas some others were only present in Hybrid: the *afaACDE* and *draP* encoding Afa/Dr adhesins, the *daaF* gene encoding the F1845 fimbrial adhesin, the *celB* gene encoding a major carbohydrate active-transport system and the *mcbA* gene encoding a bacteriocin.

**Table 3.**
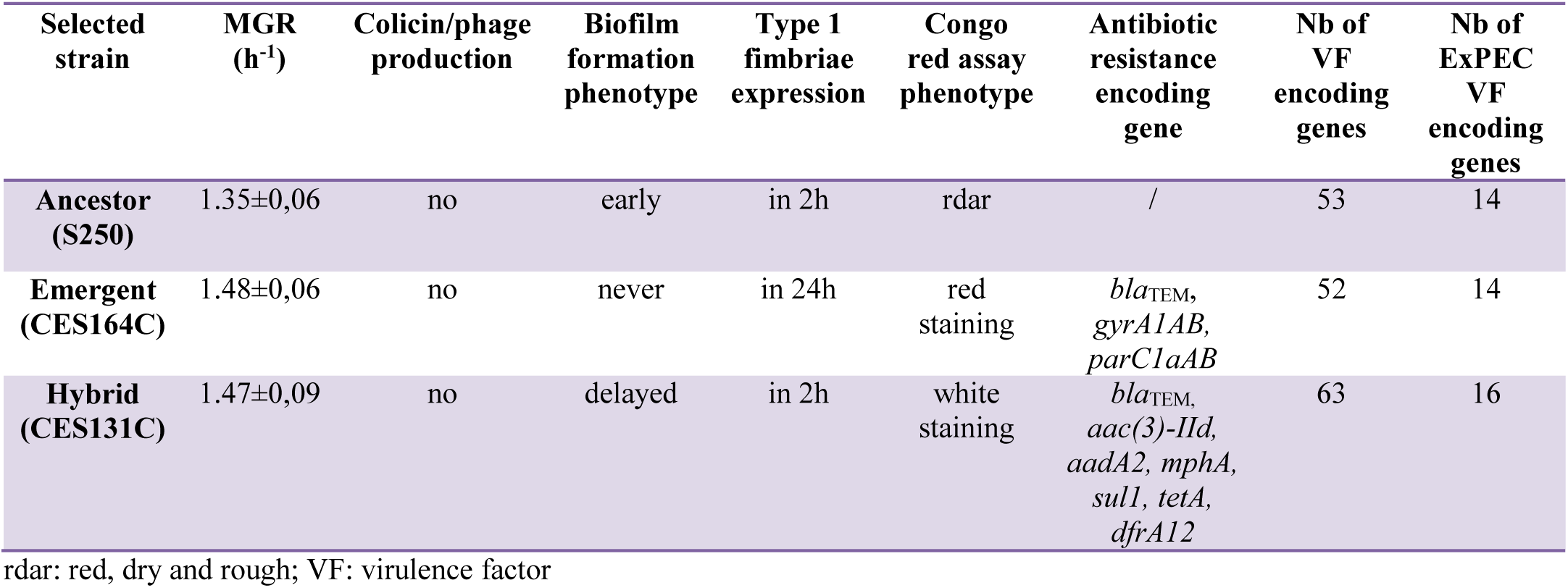
Comparative characteristics of the three strains selected for competition assays.

**Figure 4.**
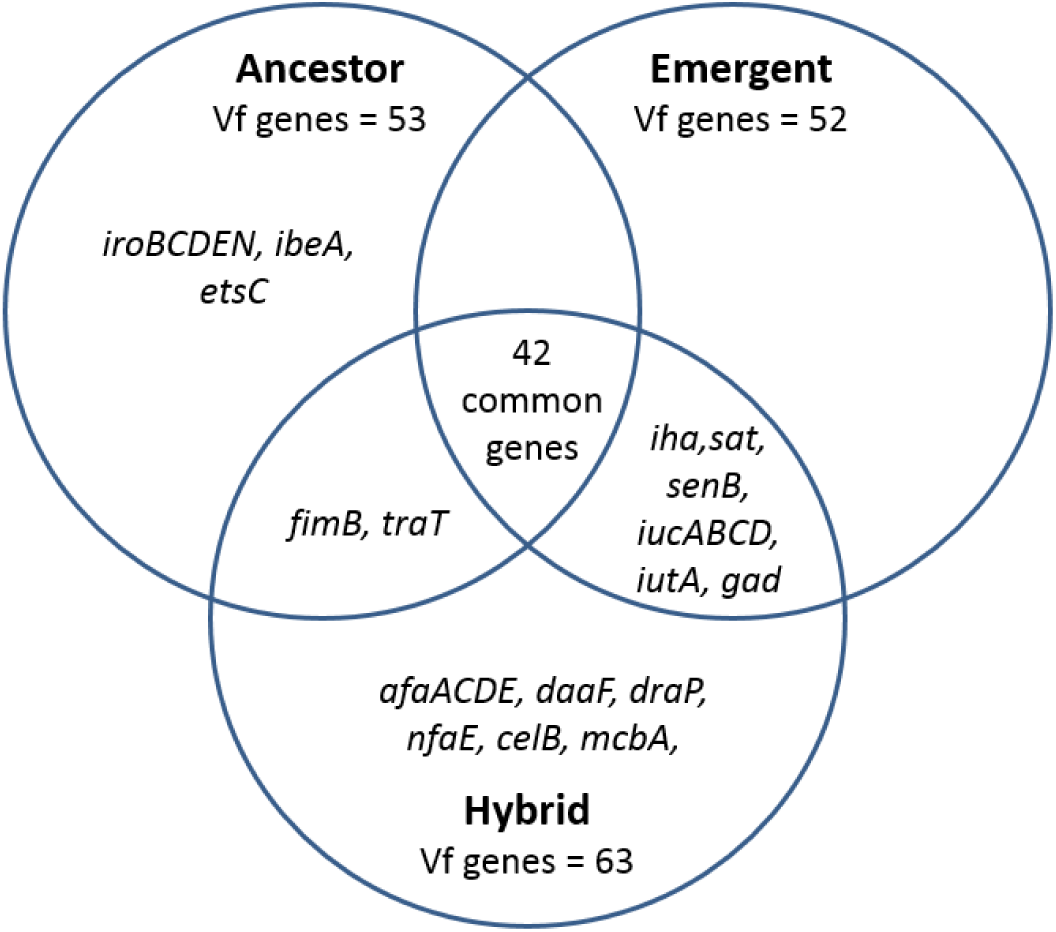
Virulence factors (VF)-encoding genes in Ancestor, Emergent and Hybrid. Each strain is represented by a circle, VF encoding genes specific of a strain are indicated in the unshared part of the circle, while genes in common are indicated in the intersecting region between the two or three involved strains. Among the 67 genes found at least once in these three strains, the 42 common genes are as follows: *fdeC, fimACDEFGHI, yfcV, yagVWXYZ, ykgK, chuA, fyuA, entABCEFS, fepABCDG, fes, irp2, kpsDEMMTII, K5, iss, malX, usp, ompT, ompA, aslA*.

To assess the potential role of the *fimB* gene in the fitness of our strains, we also retained Δ*fimB*::Kana variants of Ancestor and Hybrid, the absence of kanamycin cassette fitness cost having been checked *in vivo* by competing AncestorΔ*fimB*::Kana against AncestorΔ*fimB*:: and HybridΔ*fimB*::Kana against HybridΔ*fimB*:: (see Figures 6 and 8). We also checked the absence of fitness cost of the Δ*aada*2::Kana variant by competing HybridΔ*aada*2::Kana against Hybrid in planktonic conditions in LB, as we were obliged to use only streptomycin susceptible strains in the intestinal colonization model (data not shown).

### Competition assays

#### In planktonic conditions

No significant difference was observed between the relative fitness of Ancestor, Emergent and Hybrid in aerobic batch LB growth as CIs did not exceed 1 or −1 log (Figure 5A). The absence of the *fimB* gene did not impact the relative fitness of Ancestor and Hybrid (Figure 5A).

**Figure 5.**
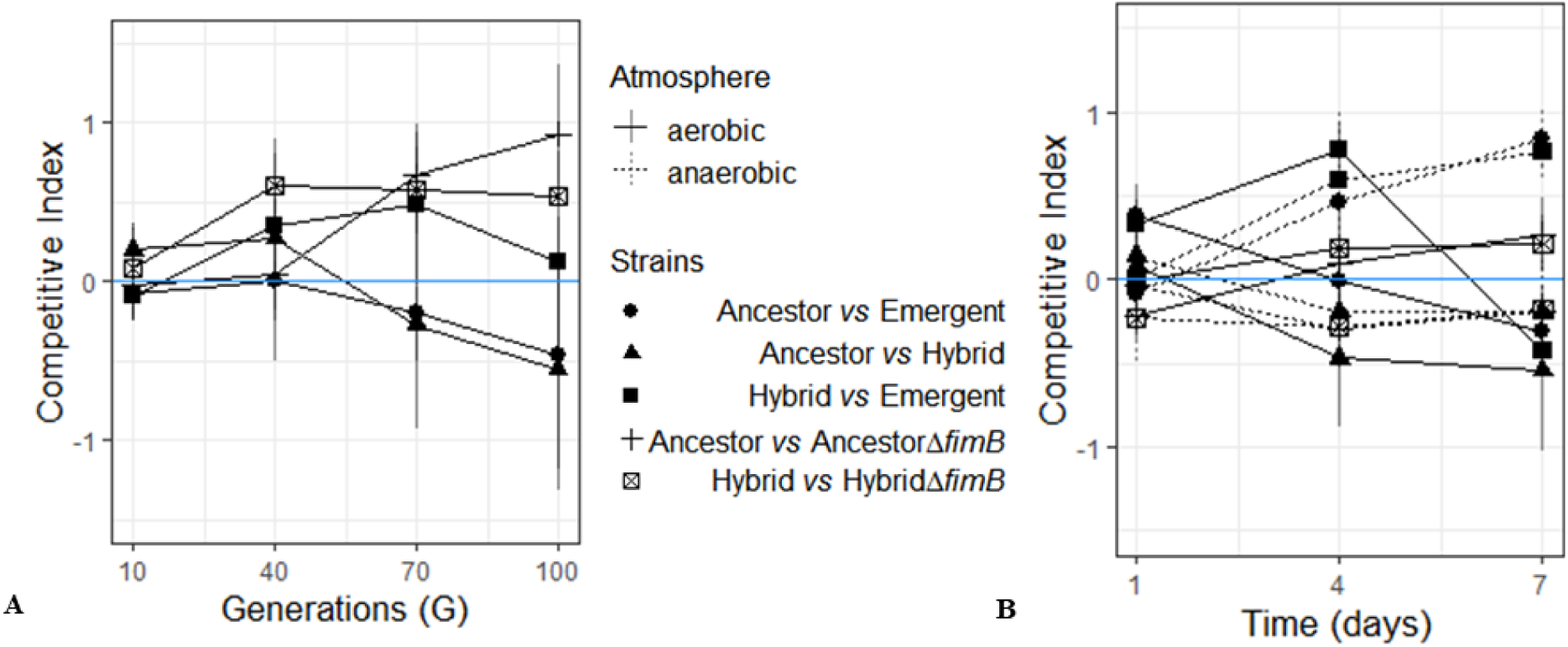
*In vitro* competitions. Dots shapes indicate the couple of strains in competition in panels A and B. Competitive index (CI) is expressed in log, and CI above zero (blue line) means that isolate 1 outcompetes isolate 2, and conversely. An interval of ±1 log was considered as the limit of discrimination. Error bars represent the standard error of the mean of a least three experiments. A. Competitions at an initial ratio of 1:1 in planktonic conditions in LB medium during 100 generations. B. Competitions at an initial ratio of 1:1 in mature biofilm conditions (bacteria spotted on BHI agar plate) during 7 days, in aerobic (continuous line) and anaerobic atmosphere (dashed line).

#### In mature biofilm conditions

No significant difference was observed between the relative fitness of Ancestor, Emergent and Hybrid in aerobic and anaerobic biofilm growth conditions as CIs did not exceed 1 or −1 log (Figure 5B). However, a small competitive advantage was observed for both Ancestor and Hybrid against Emergent, after 7 days in anaerobic biofilm conditions. Lack of the *fimB* gene did not impact the relative fitness of Ancestor and Hybrid in these conditions (Figure 5B).

#### Intestinal colonization mouse model

Ancestor displayed a better intestinal colonization capacity than Emergent. Following the streptomycin treatment, none of the mice used in this experiment had Enterobacteriaceae in feces at the time of tested bacteria inoculation. Feces sampled at each studied time contained between 10^5^ and 10^10^ *E. coli* per gram showing a satisfactory colonization rate of mice (data not shown). Ancestor outcompeted significantly Emergent after seven days post-inoculation by at least 1.5 log CFU (P = 0.02) (Figure 6), whereas there were no significant differences between Ancestor and Hybrid, and Emergent and Hybrid. Lack of the *fimB* gene did not seem to impact the colonization ability of Ancestor and Hybrid (Figure 6).

**Figure 6.**
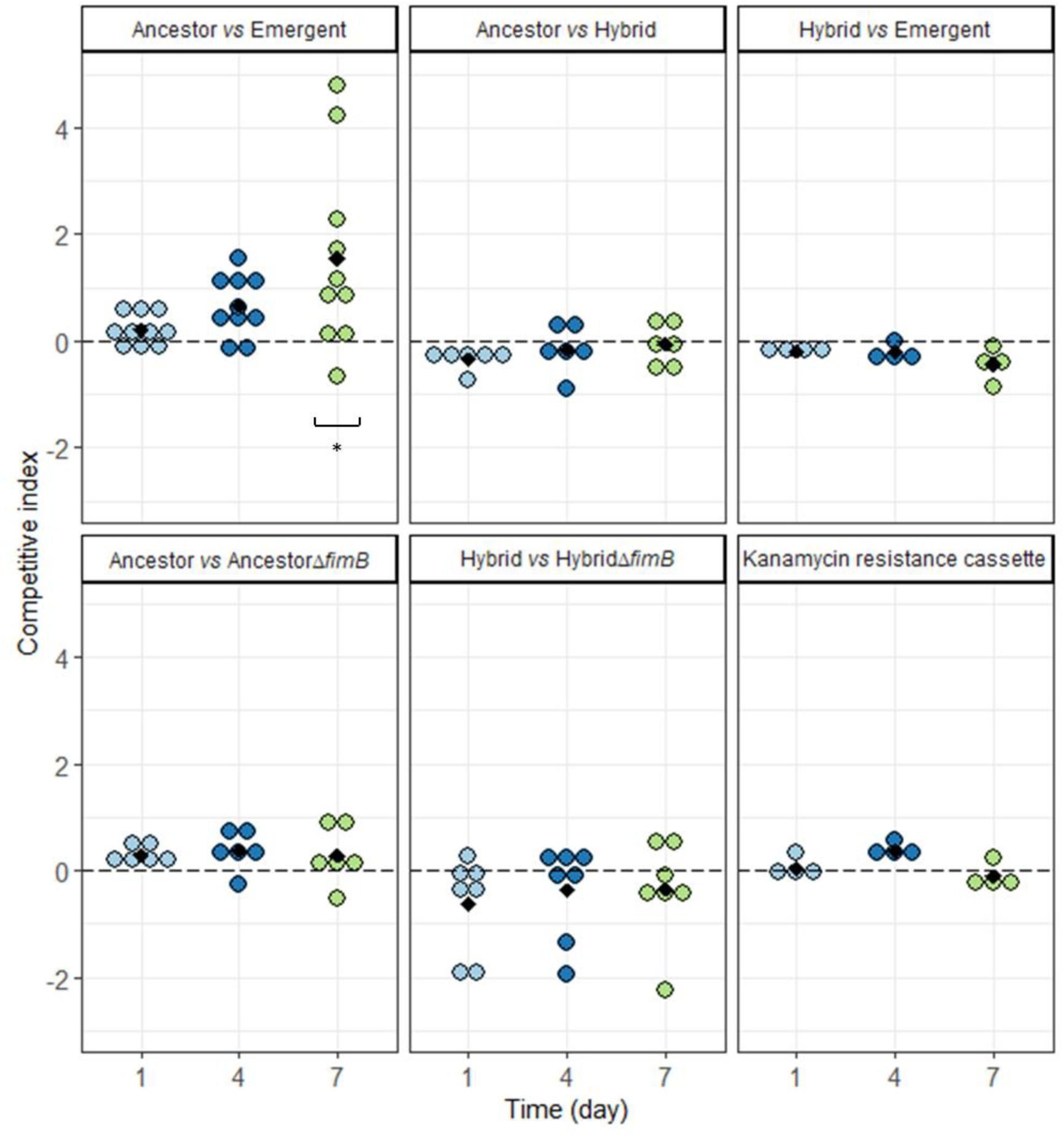
Competition in an intestinal colonization mouse model. Competitive index (CI) is expressed in log, and CI above zero means that isolate 1 outcompetes isolate 2, and conversely. An interval of ±1 log was considered as the limit of discrimination. Each dot represents a mouse at given times: light blue: day 1, dark blue: day 4 and green: day 7 post-inoculation. Black diamonds depict mean values of CI. Kanamycin resistance cassette: control competition evaluating the cost of the Kanamycin resistance cassette, with AncestorΔ*fimB*:: or HybridΔ*fimB*:: as isolate 1, and AncestorΔ*fimB*::Kana or HybridΔ*fimB*::Kana as isolate 2, respectively. * P=0.02, generated by the Wilcoxon signed-rank test and corrected for multiple comparisons by the Benjamini-Hochberg procedure.

#### Septicemia mouse model

In terms of *in vivo* virulence expression, Ancestor, Emergent and Hybrid were “killers”, as they killed between 10 and 80% of the mice 24 h after inoculation (39). Kaplan-Meier survival curves showed significantly different killing patterns (Figure 7): Ancestor killed 70% of the mice in 24 h and 100% in 26 h whereas Emergent and Hybrid, which killed 0 and 20% of the mice in 24 h, respectively, required four days to kill 100% of the mice (P<0.001). No significant difference was observed between the positive control CFT073 killing pattern and the Ancestor’s one. The lack of the *fimB* gene in Ancestor and Hybrid did not impact their killing pattern (Figure 7). In order to observe subtler differences in the *in vivo* virulence, competitions in a 1:1 ratio were performed in this mouse model. Spleen bacterial analyses showed that Ancestor outcompeted by 1 and 2 log Hybrid and Emergent, respectively (P=0.002), and Hybrid outcompeted Emergent by 1 log (P=0.002) (Figure 8). Lack of the *fimB* gene in Ancestor and Hybrid did not impact their *in vivo* virulence in this mouse model (Figure 8).

**Figure 7.**
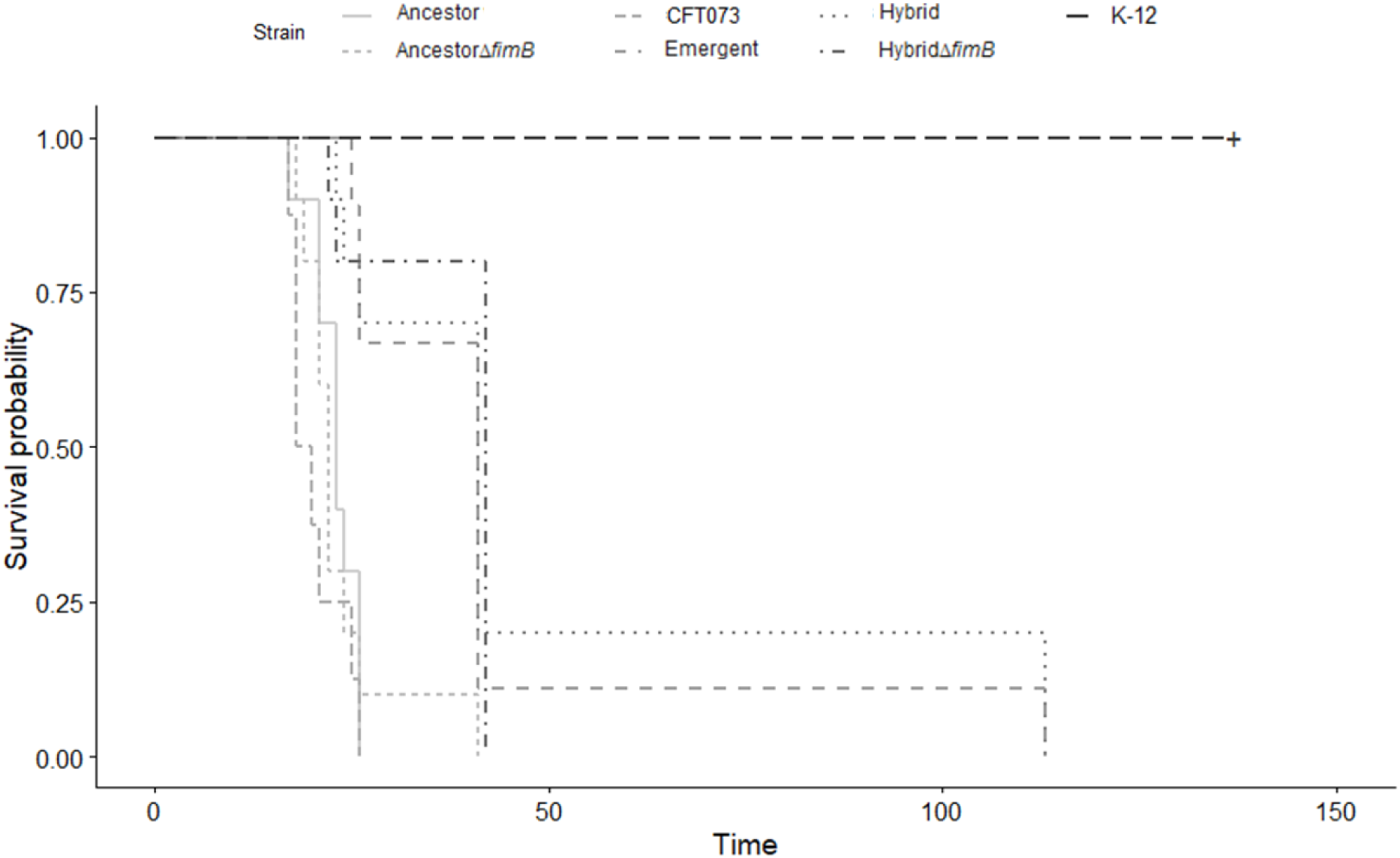
Kaplan-Meier survival curves of mice injected with *E. coli* ST131 strains. Strains are represented by line types.

**Figure 8.**
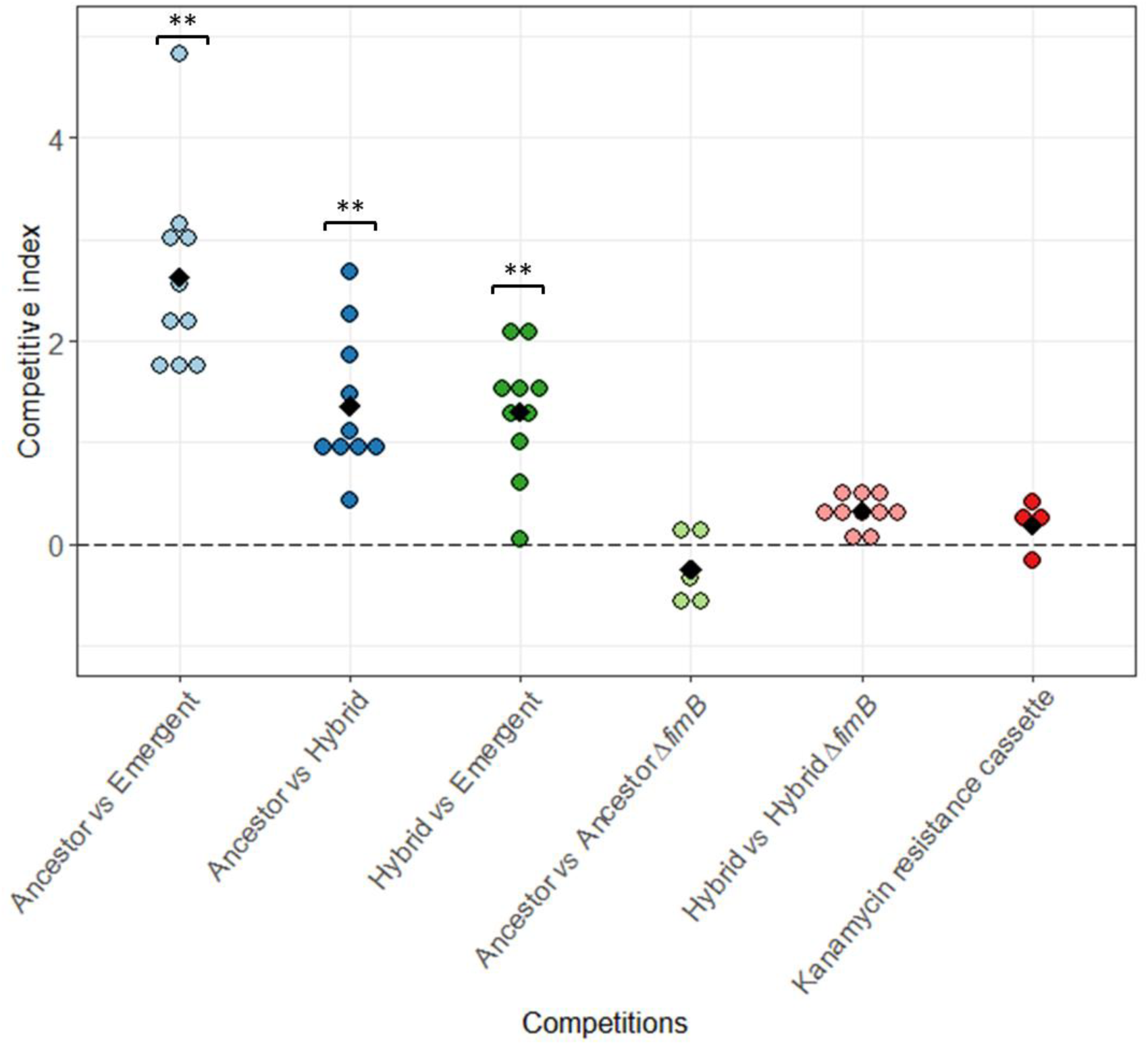
Competition in septicemia mouse model determined by spleen bacterial load. Competitive index (CI) is expressed in log, and CI above zero means that isolate 1 outcompetes isolate 2, and conversely. An interval of ±1 log was considered as the limit of discrimination. Kanamycin resistance cassette: control competition evaluating the cost of the Kanamycin resistance cassette, with AncestorΔ*fimB*:: or HybridΔ*fimB*:: as isolate 1, and AncestorΔ*fimB*::Kana or HybridΔ*fimB*::Kana as isolate 2 respectively. ** P=0.002, generated by the Wilcoxon signed-rank test.

#### Urinary tract infection mouse model

Ancestor displayed a better urinary tract colonization capacity than Emergent. In total, 30 mice were inoculated, with an overall infection rate of 87%, and no death was observed 48 h after the inoculation. Bladders and kidneys collected 48 h after inoculation contained an average of 9.10^7^ CFU/g (range: 4.10^4^-2.10^9^ CFU/g) and of 5.10^5^ CFU/g (range: 6.10^1^-2.10^6^ CFU/g), respectively, showing a satisfactory colonization rate of mice (data not shown). Ancestor outcompeted Emergent by about 2 log in the bladder and the kidneys (P=0.004) (Figure 9). Hybrid outcompeted Emergent by about 2.5 log in the bladder (P=0.008) but not in the kidneys (Figure 8). No significant difference was observed between Ancestor and Hybrid, even though Hybrid was predominant of about 2 log more in average in the kidneys (Figure 9).

**Figure 9.**
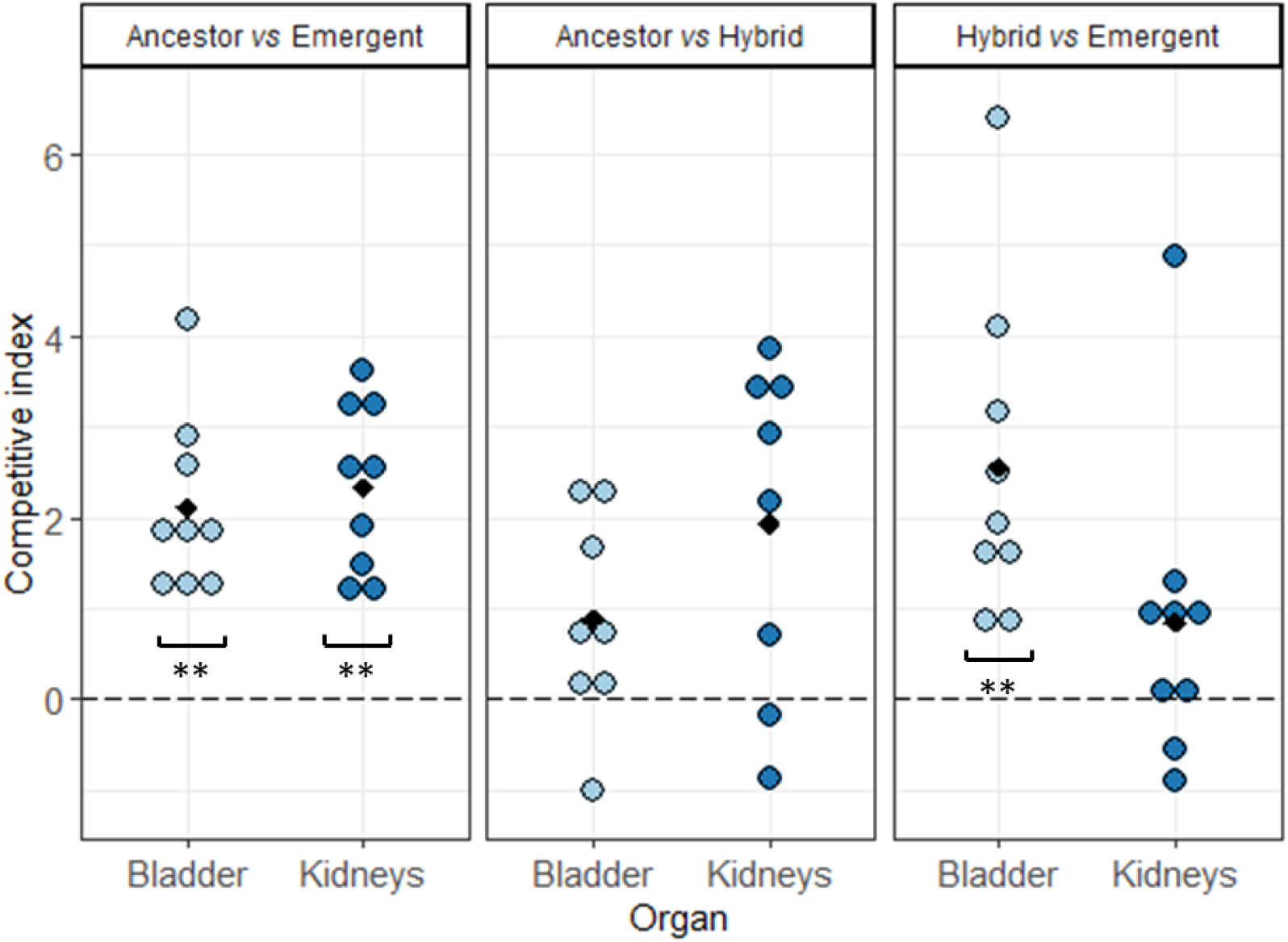
Competition in urinary tract infection mouse model determined by bladder and kidney bacterial load. Competitive index (CI) is expressed in log, and CI above zero means that isolate 1 outcompetes isolate 2, and conversely. An interval of ±1 log was considered as the limit of discrimination. ** P<0.01, generated by the Wilcoxon signed-rank test and corrected for multiple comparisons by the Benjamini-Hochberg procedure.

## Discussion

The purpose of this work was to examine the phenotypic differences between clade B and clade C strains regarding growth rate, early biofilm formation, colonization ability and virulence, in order to estimate their potential role in the success of ST131 clade C strains. The originality and strength of our study was to analyze these phenotypic differences in various i*n vitro* and *in vivo* models and under competition conditions. The limited number of strains submitted to *in vivo* competitions was determined by ethical concerns related to animal experimentations. Transposition of our *in vivo* results to human has to be made with caution because the animal models do not strictly reflect the infection physiological process observed in humans, particularly for the UTI model in which bacteria urethra ascent is absent.

Our ST131 collection comprised representative strains of subclade B and C, except for one, called Hybrid, which belonged to subclade B4 and harbored a *fimH*30 allele. This exceptional feature has already been reported in another subclade B4 strain (5), indicating that *fim* operon recombination that shaped the clade C evolution had also occurred among clade B strains, but apparently with a lower adaptive success in the genetic background of clade B than in that of clade C.

No growth rate difference was detected between our clade B and clade C strains and between fluoroquinolone susceptible and resistant strains. The latter finding is consistent with the fact that, among the mutations leading to fluoroquinolone-resistance, those observed in ST131 displayed the lowest fitness cost (11). Moreover, no growth rate difference was observed between clinical and commensal strains of our collection. This might suggest that ST131 is adapted to both commensalism and extraintestinal virulence, supporting the idea that virulence could be a derivative of commensalism (29).

The genetic and phenotypic variability of ST131 subclades were rarely explored in details in previous literature. Our analyses of Kallonen *et al.*’s blood *E. coli* isolates showed that C1 and C2 subclade strains increased in frequency over time and that clade B strains, notably those of B4 and B5 subclades, persisted as well, representing 16.5% of the ST131 isolates. One of the most interesting findings of our study was that clade C strains were less virulent and had lower intestinal and urinary colonization capacities than clade B strains. Regarding intestinal colonization, no comparison exists in the literature between these two clades, but contradictory results were obtained considering intestinal cell adhesion capacity (40). In terms of virulence, conflicting results were published, as, on the one hand, fluoroquinolone susceptible ST131 strains were shown to kill 50% of the mice when resistant ST131 strains killed 20% (41), and on the other hand, strains of virotype D, *i.e.* clade B-specific virotype, were shown to be less virulent than strains of other virotypes associated with clade C (39). Regarding urinary colonization, a clade C ST131 strain compared with strains belonging to well-known UPEC STs in separate mouse urinary tract infection (UTI) models, was shown to have a higher UTI ability (28). Here, we demonstrated that a representative subclade B1 strain (Ancestor) outcompeted a representative clade C strain (Emergent) in a UTI model. Moreover, the “Hybrid” B4 subclade strain (harboring the *fimH*30 allele) outcompeted the clade C strain in the bladder but not in the kidneys. Interestingly, Hybrid strain also displayed intermediate phenotypes with regard to intestine colonization and sepsis in mice, which could illustrate the gradual loss of virulence of ST131 over its phylogenetic evolution.

As previously described, our clade B strains were more frequently early biofilm producers than clade C strains (23, 24, 42). Knocking out the *fimB* gene in clade B strains resulted in loss of both early biofilm production capacity and early type 1 fimbriae expression. It was shown that clade C ST131 mutants devoid of type 1 pili expression had a lower intestinal and urinary tract colonization abilities compared to parent strains (25, 40). We subjected Δ*fimB* clade B mutants to competition experiments with parent strains. No differences were detected neither during growth in mature biofilm conditions nor during mouse intestinal colonization and septicemia model, suggesting that early biofilm production kinetic doesn’t seem to contribute to the better colonization abilities or to the higher virulence observed in clade B strains. In the same way, *E. coli* biofilm formation capacity on abiotic surfaces was not correlated to the ability to durably colonize mouse intestine (29). However, to confirm this hypothesis, competitions between Ancestor Δ*fimB* mutant and Emergent would be required.

It may appear surprising that the successful clade C had lower virulence and lower competitive ability in the intestine and in a UTI model. We propose two explanations. First, the resistance to fluoroquinolones displayed by clade C may come at a direct cost (43). This cost would not be revealed in maximum growth rate in planktonic conditions, but would manifest as a lower colonization ability. In spite of this cost, antibiotic resistance allowed clade C to spread. Second, the lower virulence may actually confer an additional evolutionary advantage to clade C. According to the trade-off theory (44), host exploitation by a pathogen evolves to an optimal level under a balance between the benefits in terms of transmission and the costs in terms of host mortality (45). Thus, lower virulence could confer improved fitness overall by avoiding symptomatic infections (46, 47), even if it comes at the cost of lower colonisation ability.

To conclude, we revealed substantial phenotypic differences in different clades of ST131, with the successful and multi-resistant clade C being notably less able to form biofilm, and less able to colonize the intestine and the urinary tract than its ancestor clade B. Whether these phenotypic differences represent a cost in spite of which clade C is successful, or an additional adaptation, remains an open question that would be interesting to explore in further work.

## Acknowledgments

We thank Christophe Beloin for his advices about biofilm exploration. We thank Meril Massot, Marie Vigan and Cedric Laouénan for their help with statistical analyses. We also thank Olivier Tenaillon and Antoine Bridier-Nahmias for genomic analyses assistance. Finally, we thank Johann Beghain, Mélanie Magnan and Françoise Chau for their technical assistance in this work.

This study was supported by a grant from the project JPI-EC-AMR 2016 with the French Agence Nationale de la Recherche (ANR) as sponsor (N°. ANR-16-JPEC-0002-04) to MHNC and by the “Fondation pour la Recherche Médicale” (Equipe FRM 2016, grant number DEQ20161136698) to ED. S.C.F.S. acknowledges the FPU programme for her grant (FPU15/02644) from the Secretaría General de Universidades, Spanish Ministerio de Educación, Cultura y Deporte.

## Supplemental material legend

Text S1. Technical details of the Materials and Methods section

Table S1. Strain collection: characteristics known at collection time

Table S2. Genome sequence quality

Table S3. Primers used in this study

Table S4. Plasmids used in this study

Table S5. Genotypic and phenotypic analyses of strain collection

Table S6. Main virulence factor (VF)-encoding genes

**Figure S1. Multiple correspondence analyses (MCA) of the 39 *E. coli* ST131 strains.** (A) The variables are as follows: the sources (fecal or clinical sample), the *fimH* allele (*H*22 or *H*30), fluoroquinolone (FQ) susceptibility (FQ_S: susceptible to FQ or FQ_R: resistant to FQ), the ESBL production (ESBL_P: ESBL producer or ESBL_N: non-ESBL producer), the biofilm production (early, delayed or never). cos2: square cosine (B) Plane projections on the Dim1-Dim2 of the variables and the strains. Strains or groups of strains are represented by colored circles: clade B in red and clade C in blue; circle size varies with the number of strain represented; variables are represented by black triangles. Locations of the three chosen strains, Ancestor, Hybrid and Emergent are pointed.

**Figure S2. Maximum likelihood phylogenetic tree.** This tree was built from non-recombinant SNPs using maximum likelihood, rooted on a clade A ST131 reference strain, SE15. Scale bar represents the average nucleotide substitutions per site. Branches are colored in accordance with subclades: B1 in yellow, B2 in pale orange, B3 in dark orange, B4 in red, B5 in pink, B0 in purple, C0 in pale blue, C1 in light green, C1-M27 in dark blue and C2 in dark green. Strains are named according to their origin : BZ for Ben Zakour *et al.*(5), K for Kallonen *et al.*(9) and D for this work.

**Figure S3. Schematic representation of the *fim* operon in a clade B strain, a clade C strains and Hybrid strain of subclade B4.** Genes are depicted by arrows except for *fimS* that is an invertible element, and the dotted lines represent ISEc55: the insertion sequence present within the *fimB* gene of all clade C strains. A. Comparison between a representative clade B strain and Hybrid . B. Comparison between Hybrid and a representative clade C strain. SNP: single nucleotide polymorphisms and their effect: non-synonymous and synonymous. The additional number indicates the presence of gap.

